# Contrasting levels of transcriptome-wide SNP diversity and decoupled patterns of molecular and functional adaptive variation in conifers

**DOI:** 10.1101/2023.12.12.571309

**Authors:** Nathalie Pavy, Sébastien Gérardi, Julien Prunier, Philippe Rigault, Jérôme Laroche, Gaétan Daigle, Brian Boyle, John Mackay, Jean Bousquet

## Abstract

Adaptive convergence can arise when response to natural selection involves shared molecular or functional mechanisms among multiple taxa. Conifers are of ancient origin with delayed sexual maturity related to their woody perennial nature. Thus, they represent a relevant plant group to assess if convergence from selection may have become disconnected between molecular and functional levels. In this purpose, transcriptome-wide SNP diversity was assessed in seven partially sympatric and reproductively isolated conifer species populating the temperate and boreal forests of northeastern North America. SNP diversity was found highly heterogeneous among species, which would relate to variation in species-specific demography and history. Rapidly evolving genes with signatures of positive selection were identified, and their relative abundance among species reflected differences in transcriptome-wide SNP diversity. Their analysis also revealed very limited convergence among taxa in spite of sampling same tissues at same age. However, convergence increased gradually at the levels of gene families and biological processes, which were largely related to stress response and regulatory mechanisms in all species. Given their multiple small to large gene families and long time since inception, conifers may have had sufficient gene network flexibility and gene functional redundancy for evolving alternative adaptive genes for similar metabolic responses to environmental selection pressures. Despite a long divergence time of ∼350 Mya between conifers and Angiosperms, we also uncovered a set of 20 key genes presumably under positive selection in both lineages.

## INTRODUCTION

Adaptive genetic variation allows organisms to cope with natural selective pressures and thrive in their environment. This is especially true for long-lived woody plants, such as conifers from mid-northern latitudes, that must contend with delayed sexual maturity to adapt to highly heterogeneous and changing climatic conditions (Depardieu et al., 2021). Therefore, identifying and characterizing adaptive genetic variation within species is crucial to understand the molecular mechanisms underlying their response to environmental pressures. Molecular convergence can arise when such molecular mechanisms are shared by multiple species (Stern, 2013). This process may occur at different hierarchical levels, such as specific nucleotides, protein-coding genes (often referred to as ‘gene reuse’), gene families, or genes belonging to the same biological pathways (Hao et al., 2019; Sackton & Clark, 2019). As a general trend, molecular convergence is expected to increase with hierarchical levels under similar positive selection pressures (Stern, 2013; He et al., 2020; Xu et al., 2020) and thus, it is expected that significant divergence in adaptive evolution must exist at the gene level.

However, the many determinants of molecular convergence complicate the prediction of patterns of adaptive evolution at both intraspecific and interspecific taxonomic levels. The most influential determinants include ancestry (the probability of convergence decreases along with taxa divergence time), effective population size (taxa of small effective population size are less likely to converge due to increased genetic drift), gene flow / introgression (gene flow usually increases convergence by constraining differentiation among taxa, but can also prevent local adaptation), selection landscape (convergence is expected to decrease when the number of selective pressures increases in a given habitat), and many-to-one mapping (convergence is expected to decrease as the number of traits governing a given functional output increases) (reviewed by Bolnick et al., 2018).

Many-to-one mapping is a particularly relevant determinant of molecular convergence when studying the adaptive trajectories of species. Indeed, considering that the link between phenotypic and molecular convergence is well established in a variety of taxa (see Martin & Orgogozo, 2013 for a catalog of genetic hotspots of phenotypic variation in animals, plants, and yeast), it is reasonable to assume that molecular convergence reflects shared adaptive response to similar selective pressures. However, the opposite is not necessarily true. The fact that adaptative traits are usually highly polygenic (e.g. Le Corre & Kremer, 2012; Csilléry et al., 2018; Barghi et al., 2020; Depardieu et al., 2021) suggests that plant taxa have typically many genetic solutions available to solve the adaptive challenges they face in nature (Arendt & Reznick, 2008; Losos, 2011; Tenaillon et al., 2012; Storz, 2016), including in closely-related populations from the same species (e.g. Manceau et al. 2010; Elmer & Meyer, 2011). Most plant groups such as conifers are also characterized by large gene families and redundancy of gene function (Guillet-Claude et al., 2004; Bedon et al., 2010; Pavy et al., 2012a; Stival Sena et al., 2018; Van Ghelder et al., 2019). Therefore, species may follow similar adaptive trajectories, while showing reduced levels convergence at the molecular level. Hence, assessing the extent of functional convergence of adaptive genes in multiple species can complement the picture derived from molecular convergence alone, and reveal otherwise hidden adaptive patterns.

To address these fundamental questions related to adaptive convergence, conifers from northeastern North America represent an ideal framework for several reasons. First, contrary to European forests for instance, these forests have been generally characterized by low levels of anthropic disturbance up to the twentieth century (i.e. reduced urbanization and forest management) and the regional landscape is of relative topographic homogeneity, compared to western North America for instance. With the limited potential of confounding factors from long-term human activity and the lack of significant barriers constraining migration during the Holocene, tree species could track their most suitable current habitats and evolve local adaptations in response to environmental selective pressures, as evidenced by several empirical studies (e.g. Namroud et al., 2010; Prunier et al., 2011; Hornoy et al., 2015; Nadeau et al., 2016). Second, none of the conifers in the boreal forest of northeastern North America are known to hybridize, although they are sympatric in most of their range: potential hybrid zones are all located at the southern or western edge of the species ranges, they have been quite well delimited and are therefore easy to avoid by using an adequate sampling strategy (Jaramillo-Correa et al., 2009), which would minimize the risk that molecular signatures of natural selection within species are confounded by interspecific introgression. Also, the extensive range overlap of conifer species across the mid-latitude forests of northeastern North America indicates that these species generally face common environmental pressures, of which harsh and heterogenous climatic conditions are a large component (Hornoy et al., 2015; Depardieu et al., 2021). Thus, these conifers represent relevant models to address questions about long-term evolution and adaptation from a comparative perspective.

However, studying molecular convergence in woody perennial plant taxa with such large and complex genomes is highly challenging. Over the last 20 years, our understanding of conifer genomes has progressed significantly through the sequencing and analyses of their genome structure, evolution and functions (reviewed by Prunier et al., 2016). Nonetheless, extensive resequencing has been restricted to only a few conifer species belonging primarily to the *Picea* and *Pinus* genera. This limits the potential to conduct exhaustive comparative studies between species, which are essential to understand the common determinants of adaptive evolution. To date, the main findings indicate a rather limited convergence among adaptive genes identified from species belonging to the same or different genera (Mosca et al., 2012; Yeaman et al., 2016; Bousquet et al., 2021; Gagalova et al., 2022).

In this study, we investigated adaptive convergence among six sympatric Pinaceae species native of northeastern North America, namely white spruce (*Picea glauca*), black spruce (*Picea mariana*), eastern white pine (*Pinus strobus*), jack pine (*Pinus banksiana*), balsam fir (*Abies balsamea*), and tamarack (*Larix laricina*), as well as one sympatric Cupressaceae taxon, eastern white cedar (*Thuja occidentalis*). Considering that the Cupressaceae and the Pinaceae diverged ∼315 Mya (Leslie et al., 2018), while taxa divergence within the Pinaceae did not take place before ∼185 Mya (divergence of the *Abies* genus from its sister taxa; Leslie et al., 2018), we included a Cupressaceae taxon to qualitatively assess the effect of phylogenetic relatedness on our inferences. In this study, we first identified gene nucleotide polymorphisms within each species, and assessed their level of overall genetic diversity across the transcriptome. We then identified rapidly-evolving genes with signatures of positive selection in each species, in order to estimate the level of adaptive convergence among species from a molecular and functional perspectives. This approach also allowed us to identify shared drivers of adaptive evolution among species. We also investigated the extent of adaptive molecular convergence between Angiosperms and this group of conifers.

## RESULTS

### SNP diversity

This study enabled the identification of almost 1.5 million of SNPs across the transcriptomes of the seven conifers analyzed (*Picea glauca* and *Picea mariana*, *Pinus strobus* and *Pinus banksiana*, *Larix laricina*, *Abies balsamea*, *Thuya occidentalis*). However, we only retained the ∼866K SNPs with highest quality to conduct the subsequent analyses. Among them, ∼398K quality SNPs were located in coding sequences representing almost ∼97K Open Reading Frames (∼16K transcripts per species, on average) (Table 1). All species considered, ∼82% of the transcripts and ∼70% of the ORFs carried SNPs. In such transcriptome sequencing project, it appeared important to minimize the effects caused by read depth before undertaking any analysis of SNP diversity. We carefully adjusted the SNP diversity by both sequence length and depth upstream of data comparison across genes and across species (see Methods). After these adjustments, overall SNP diversity was estimated for each species and it was found significantly heterogeneous among the seven conifer taxa: three groups of overall SNP diversity were delineated based on the results of Kolmogorov-Smirnov and Cramer-von Mises tests (Figure 1; Table 1; Methods S7). The group of species with the highest level of overall SNP diversity included the two *Picea* species, the group with the lowest diversity included *Pinus strobus* and *Thuja occidentalis*, while the three remaining species, *Abies balsamea, Larix laricina,* and *Pinus banksiana*, had intermediate overall SNP diversity (Table 1; Table S1). In addition, a Kruskal-Wallis test revealed significant differences in rates of synonymous, nonsynonymous and total SNPs across the seven species (Table S2).

**Figure 1.**
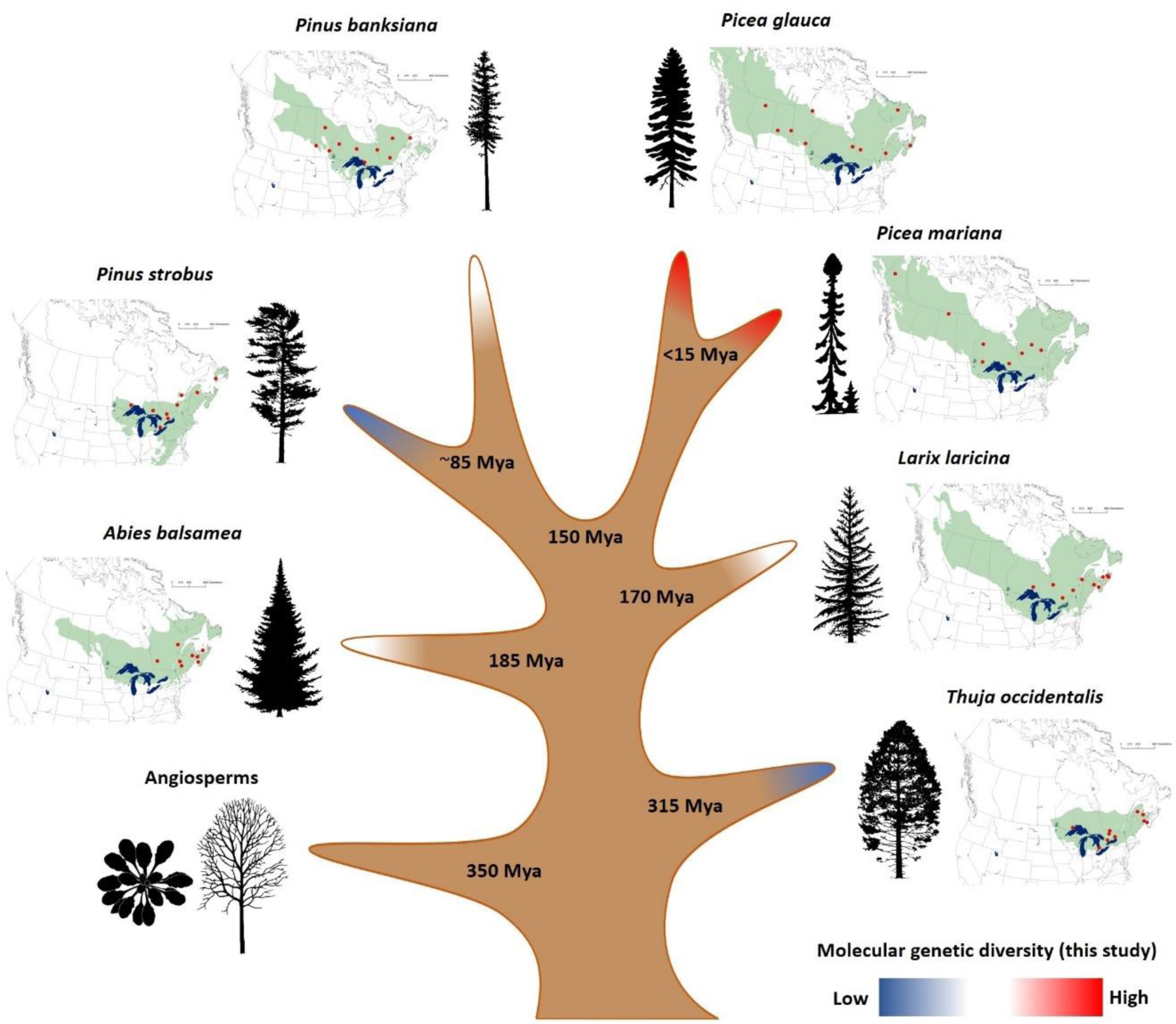
Characteristics of the seven conifer species analyzed in this study. The approximate natural range of each species is represented in green besides its associated tree silhouette, and populations sampled are mapped as red dots. For illustrative purposes only, the figure shows species divergence times derived from fossil-calibrated molecular clocks (Bouillé & Bousquet, 2005; Leslie et al., 2018; Li et al., 2019). Colors at branch tips represent the three groups of overall SNP diversity identified in this study at the transcriptome-wide level (see Results). Source of tree silhouettes: https://tidcf.nrcan.gc.ca/; https://www.phylopic.org/

**Table 1.** Metrics about high-quality SNPs for the seven conifer transcriptome datasets, including adjusted metrics for variations in sequence length and read coverage. In column 1, species were grouped according to their level of intraspecific molecular genetic diversity, based on statistical tests performed on adjusted SNP diversity data (Methods S7).

| Overall SNP<br>diversity<br>group | Species | Raw<br>number of<br>transcripts | Raw<br>number<br>of SNPs | Adjusted<br>number of<br>SNPs | Number of<br>transcripts<br>with SNP(s)<br>after<br>adjustment | Average<br>number of<br>SNPs per<br>transcript after<br>adjustment | Proportion of<br>transcripts<br>with SNPs<br>after<br>adjustment |
| --- | --- | --- | --- | --- | --- | --- | --- |
| Highest | *Picea glauca* | 18,060 | 105,778 | 100,128 | 16,581 | 6.0 | 91.8% |
| Highest | *Picea mariana* | 20,534 | 114,771 | 108,019 | 18,558 | 5.8 | 90.4% |
| Intermediate | *Pinus banksiana* | 19,510 | 90,985 | 87,011 | 16,441 | 5.3 | 84.3% |
| Intermediate | *Abies balsamea* | 19,487 | 91,311 | 86,564 | 16,521 | 5.2 | 84.8% |
| Intermediate | *Larix laricina* | 20,950 | 94,359 | 89,877 | 16,935 | 5.3 | 80.8% |
| Lowest | *Pinus strobus* | 21,795 | 71,576 | 69,867 | 15,412 | 4.5 | 70.7% |
| Lowest | *Thuja occidentalis* | 19,543 | 64,872 | 62,749 | 13,834 | 4.5 | 70.8% |

### Detection of genes with high SNP A/S ratios and relationship with overall SNP diversity

For each species and each gene where SNPs were detected in a given species, synonymous and nonsynonymous sites were identified and SNPs were classified as synonymous and nonsynonymous so to estimate the rates of synonymous (S) and nonsynonymous SNPs (A), and calculate the gene SNP A/S ratio (see Methods). When this ratio exceeds the value of 1 expected under neutral expectations, there is an excess of nonsynonymous over synonymous SNPs, which is indicative of positive or balancing selection related to adaptive evolution (Kimura, 1983; Fay et al., 2001). Such an excess of nonsynonymous SNPs was found for many genes in all species studied herein, with SNP A/S value above 1 ranging between 13% and 16% of the genes depending on the species. Moreover, whatever the species, more than 2.5% of the genes harbored an A/S value above 2, thus well beyond 1 and indicative of a strong positive selection signature (Figure 2). The proportion of genes with such high SNP A/S values was also highly correlated with the level of overall SNP diversity detected within each species (*R^2^*= 0.87; p-value < 0.01; Figure 2).

**Figure 2.**
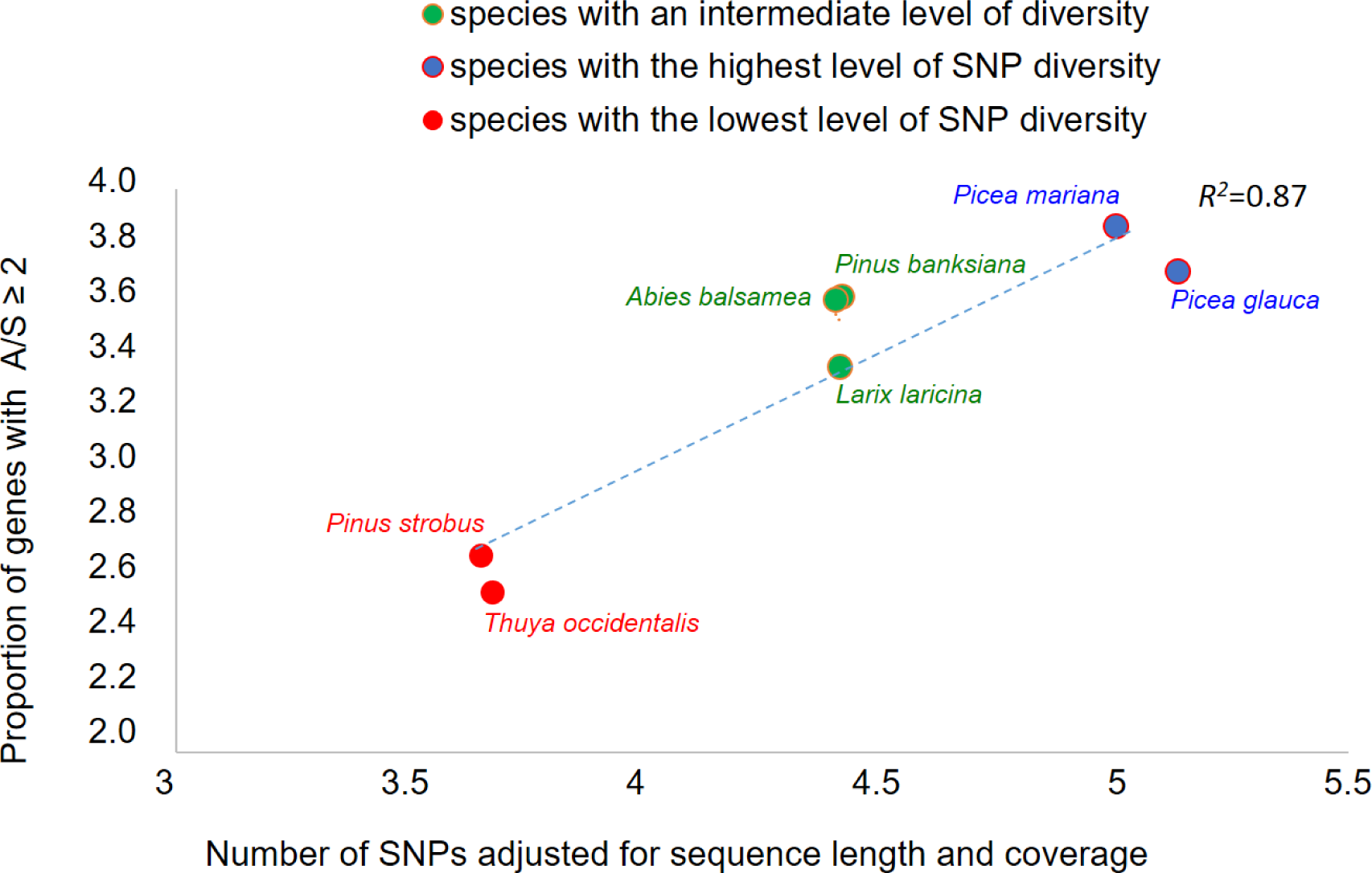
Relationship between overall SNP diversity and adaptive variation. In each species, the proportion of genes with A/S ≥ 2 was calculated as the number of genes with A/S ≥ 2 divided by the total number of ORFs in the species considered.

Also worth of mention, for each species and following increasing values of SNP A/S ratio for the genes of that species, a reverse general trend was observed between the rate of nonsynonymous SNPs versus the rate of synonymous SNPs (Figure S1). As the gene SNP A/S ratios increased, the rate of nonsynonymous SNPs increased and the rate of synonymous SNPs decreased, instead of remaining stable as expected under neutral evolution (Figure S1). Such a trend suggests the presence of purifying selection for A/S values below 1 and positive selection for A/S values above 1. Synonymous SNPs were thus essentially absent from genes with highest SNP A/S values, and this pattern was much consistent across the seven conifer species analyzed.

### Annotations of genes with the highest SNP A/S values

To ensure comparability across species regarding the detection of genes with signatures of positive selection, seven sub-datasets with identical number of such genes were delimited by identifying for each species the 300 genes with top SNP A/S values. These 300 genes represented between 2 to 3% of the genes with SNPs in each species, and these genes all harbored SNP A/S values ≥ 2.5 (Table S3). In this set of seven times 300 genes per species, thus representing a total of 2,100 genes, 67.4% of them (1,416 genes) had a significant match (blastp E-value <1E-15) with a SwissProt-Uniprot protein, a proportion consistent with other studies in conifers (Hart et al., 2020). Moreover, 1,327 genes (63.2%) had a match with 620 PFAM families (match E-value <1E-15). These genes had a wide variety of annotations (Figure S2). Among GO terms, 1,201 Biological Processes (BPs) were assigned to genes with highest SNP A/S values. The most represented processes were directly related to responses to biotic and abiotic stresses, as well as various related processes (Figure S3). For instance, five terms describing plant responses to pathogens were found 443 times. Unsurprisingly, signal transduction, which is a common denominator of cell response to a stimulus, was the most predominant term (11.6% of the annotated genes), along with defense response (9.1% of the annotated genes) (Figure S3 - Figure S4). The nicotinamide adenine dinucleotide (NAD) catabolic process was also highly represented, which is consistent with the central role of NAD in plant defense responses (Pétriacq et al., 2013).

### Diversity in gene sequences, functions and processes found among genes with highest SNP A/S values

The complete dataset of gene sequences was successfully clustered into orthogroups, indicating that there was a good overlap of transcripts sequenced across the seven conifer species (Table S4). Altogether, 94.7% of the gene sequences were assigned to an orthogroup and the remainder were orphans. Out of the 16,982 orthogroups delimited in total, 8,647 contained gene sequences from the seven species, and 1,034 others contained gene sequences from all species except the more phylogenetically distant Cupresseae taxon *Thuja occidentalis* (Figure 3A). The number of species-specific genes was low in Pinaceae taxa (between 7% and 10%) and slightly higher (15%) in *Thuja occidentalis*, which was expected given that this taxon belongs to the more divergent Cupressaceae family (Figure 3A). These species-specific gene sequences could either not be assigned to any orthogroup, or represent species-specific orthogroups (Table S4). In contrast to the trend observed in the complete dataset, orthogroups derived from gene sequences with highest SNP A/S values in each species showed a much lower overlap among species (Figure 3B). The vast majority of them were species-specific (total of 1,489; 76.9%), with only seven orthogroups shared across all seven species (Figure 3B; Figure 4). On average, species shared 42 to 43 orthogroups representative of genes with highest SNP A/S values (Table 2A).

**Figure 3.**
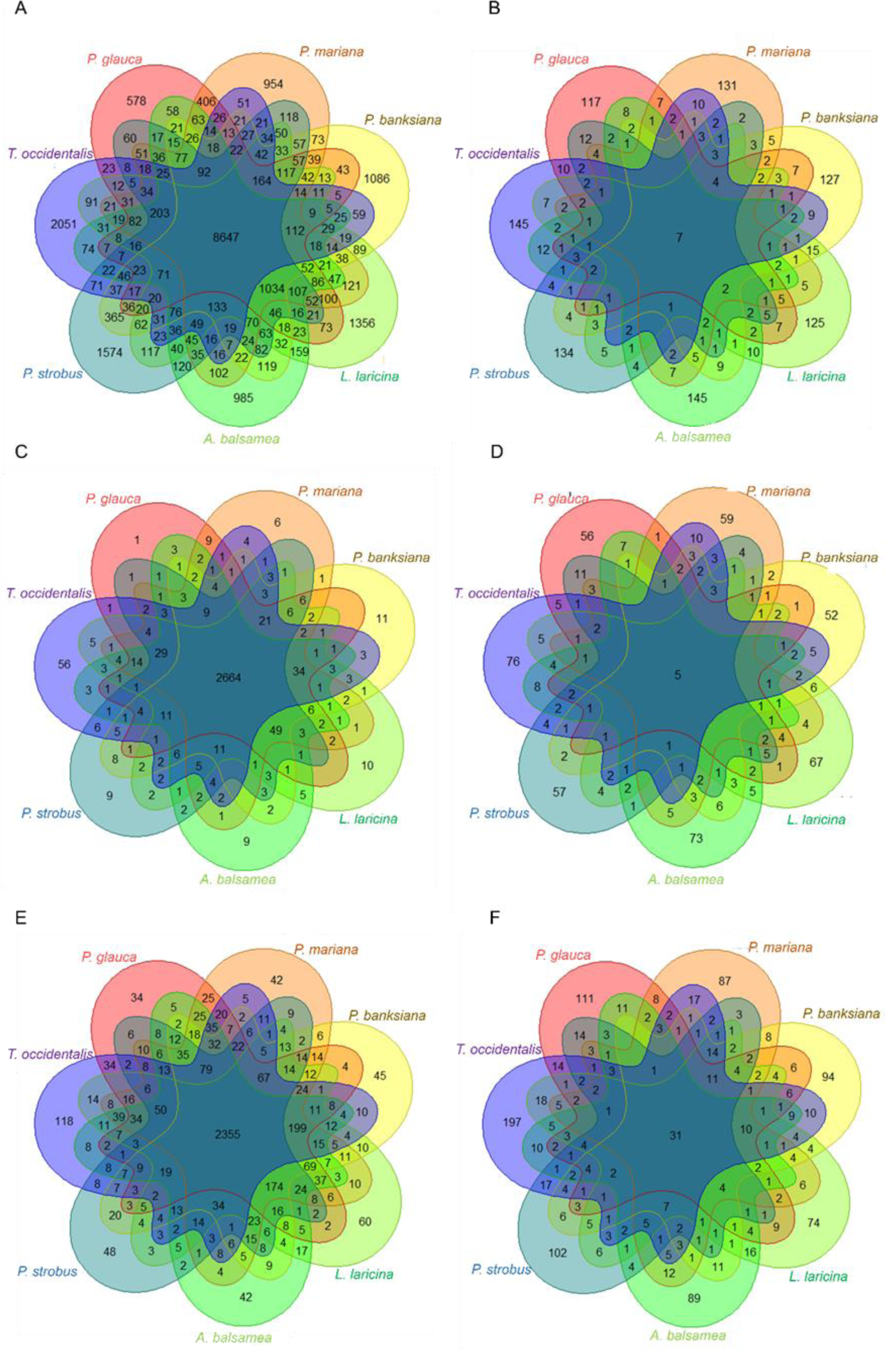
Overlap among species of orthogroups (A, B), PFAM families (C, D) and Gene Ontology Biological Processes (E, F). A, B) Orthogroups were identified by clustering the complete dataset of 139k gene sequences of all seven conifer species (*Picea glauca, Picea mariana, Pinus banksiana, Abies balsamea, Larix laricina, Pinus strobus, Thuya occidentalis*) (A), and by clustering the 300 gene sequences with highest SNP A/S values in each species (B). The number reported in each intersection corresponds to the number of orthogroups shared by species, while the number reported in each species-specific area corresponds to the number of singleton sequences that either could not be assigned to any orthogroup, or represented species-specific orthogroups. C, D) Protein families were identified based on similarities against the PFAM database across the overall transcript datasets (C) and across the 300 genes with highest SNP A/S values in each species (D). E, F) Biological processes GO terms across the overall sequence dataset (E) and across the 300 genes with highest SNP A/S values in each species (F).

**Figure 4.**
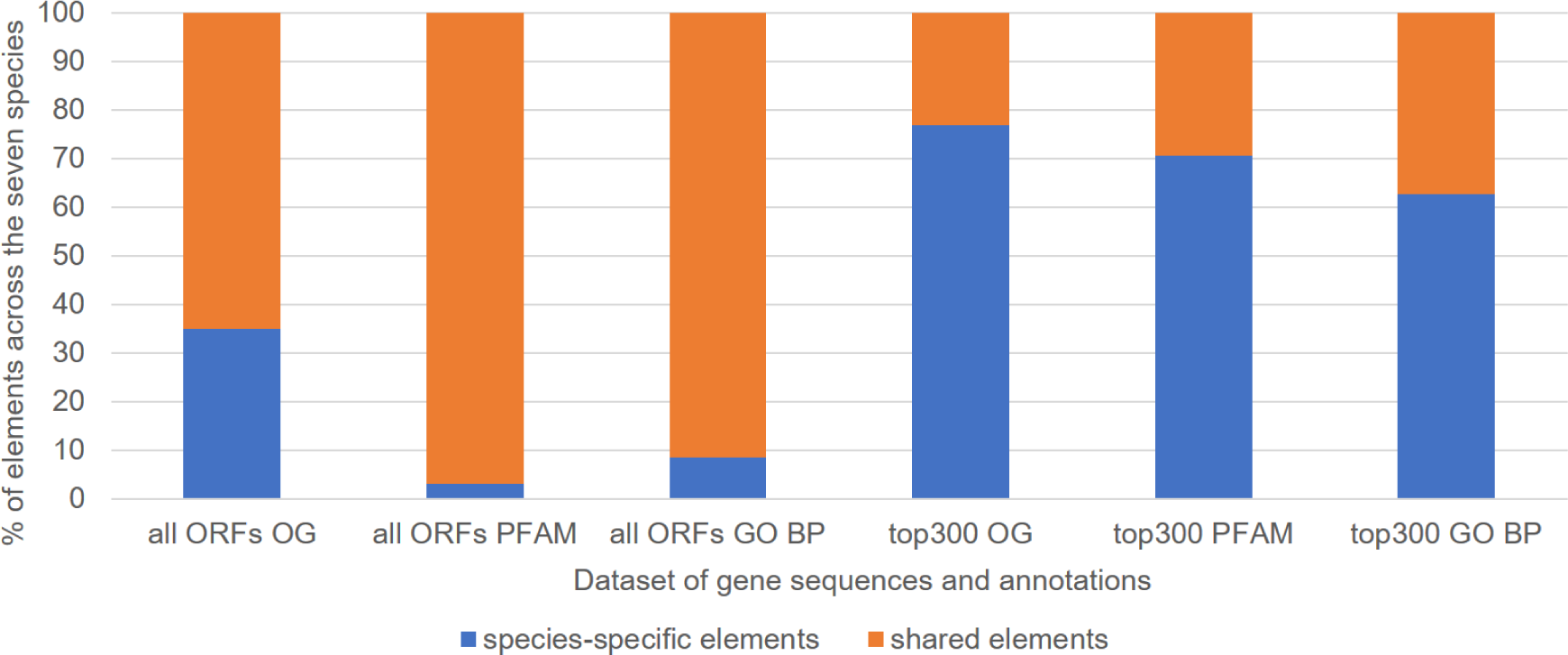
Percentage of shared or species-specific genes and annotations among the complete dataset (all ORFs) and among the 300 genes with highest A/S values in each species (top 300). Sequence clustering was performed using orthofinder (Emms & Kelly, 2019) and orthogroups (OG) overlap among species (i.e. shared by two or more species) was assessed. Sequence clustering was performed on the overall dataset of protein sequences (all ORFs) and on sequences with highest A/S values (top 300). Protein families were determined based on matches with domains or families from the PFAM database. For each protein, the match with the lowest E-value below E-15 was retained. Gene Ontology (GO) terms assigned to the Uniprot protein matching the conifer gene sequence with the lowest E-value below E-15 were used to illustrate the overlap in Biological Processes (BP) among species.

**Table 2.** Shared orthogroups and biological processes (BPs) associated with genes with highest A/S values between each pair of species. PG: white spruce (*Picea glauca*), PM: black spruce (*Picea mariana*), PB: jack pine (*Pinus banksiana*), AB: balsam fir (*Abies balsamea*), LL: tamarack (*Larix laricina*), PS: eastern white pine (*Pinus strobus*), TO: eastern white cedar (*Thuja occidentalis*).

| Family | Taxa | PG | PM | PB | AB | LL | PS | TO | Average number of shared orthogroups |
| --- | --- | --- | --- | --- | --- | --- | --- | --- | --- |
| Pinaceae | PG |  |  |  |  |  |  |  | 43 |
|  | PM | 43 |  |  |  |  |  |  |  |
|  | PB | 39 | 39 |  |  |  |  |  |  |
|  | AB | 38 | 34 | 41 |  |  |  |  |  |
|  | LL | 52 | 46 | 53 | 41 |  |  |  |  |
|  | PS | 54 | 37 | 42 | 34 | 46 |  |  |  |
| Cupressaceae | TO | 44 | 45 | 41 | 40 | 42 | 41 |  | 42 |

B. Number of shared Biological Processes (BPs)
| Family | Taxa | PG | PM | PB | AB | LL | PS | TO | Average number of shared BPs |
| --- | --- | --- | --- | --- | --- | --- | --- | --- | --- |
| Pinaceae | PG |  |  |  |  |  |  |  | 105 |
|  | PM | 100 |  |  |  |  |  |  |  |
|  | PB | 99 | 116 |  |  |  |  |  |  |
|  | AB | 97 | 98 | 98 |  |  |  |  |  |
|  | LL | 107 | 121 | 120 | 105 |  |  |  |  |
|  | PS | 101 | 105 | 112 | 85 | 114 |  |  |  |
| Cupressaceae | TO | 112 | 135 | 131 | 120 | 139 | 128 |  | 128 |
| Number of genes |  | 300 | 300 | 300 | 300 | 300 | 300 | 300 |  |
| Number of annotated genes (*) |  | 188 | 217 | 196 | 200 | 206 | 200 | 209 |  |
| Number of annotated genes/BP term |  | 0.60 | 0.58 | 0.63 | 0.58 | 0.64 | 0.59 | 0.47 |  |

(\*) based on matches against Uniprot (blastp E-value<E-15)

A similar picture was observed at the gene family level for the whole dataset. Only 102 PFAM accessions (3.2%) containing genes with highest SNP A/S values were species-specific in the overall dataset, indicating that protein families or domains were predominantly shared among the seven conifer species (Figure 3C). Nonetheless, while a majority of them (total of 440 PFAM accessions, 70.7%) were species-specific in genes with highest SNP A/S values (Figure 3D), the proportion of shared families between species increased as compared to that for orthogroups (Figure 4).

Among GO terms, 4,504 BPs and 1,201 BPs were associated with the overall dataset and the genes with highest SNP A/S values, respectively (Figure 3E; Figure S5). Species-specific BPs were relatively few in the overall sequence dataset (total of 389, 8.6%) and were still dominant (total of 754, 62.7%) in the set of genes with highest SNP A/S values, although at a lower rate than that observed for orthogroups of PFAM familes. In spite of more convergence observed at the level of BPs, these results are indicative of a high level of functional diversity in the various sets of genes with high SNP A/S values (Figure 3F; Figure 4).

The few genes with highest SNP A/S values shared by all the seven species (Figure 3B) were all homologous to sequences of known function. The five PFAM accessions shared by the seven species (Figure 3D) were also the most abundant among the 2,100 genes with highest SNP A/S values. They included the NB-ARC domain (PF00931; 99 sequences), the protein kinase domain (PF00069; 44 sequences) and the TIR domain (PF01582; 37 sequences), which are frequently found in combination in proteins involved in defense responses. The families with members having high SNP A/S values in all seven studied species included three disease resistance families (ROQ1, RPS5 and RUN1) (Table S5), the PPR repeat family (PF13041; 31 genes) and the cytochrome P450 family (PF00067; 25 genes). The most abundantly represented and shared families are detailed in Figure S5). At the gene ontology level, 31 BPs were shared by all species (Figure 1F), which represents approximatively a third of the average number of species-specific BPs (∼93 BPs) in the Pinaceae taxa (Figure 1F). As it was the case for genes with high SNP A/S values, the Cupressaceae *Thuja occidentalis* had the highest number of species-specific BPs (∼200 BPs; Figure 1F). While the Pinaceae taxa shared an average of 105 BPs with each other, they also shared an average of 128 BPs with *Thuja occidentalis* (Table 2B). The 31 terms shared by all conifer species could be categorized into seven groups: responses to stimuli, regulation of defense response, development, cell division, cell wall organization, metabolic processes and nucleotide metabolic process (Figure S6). More than half of the terms fell into response to stimuli (like jasmonic acid or salicylic acid) and responses to biotic (like detection of bacterium, response to fungi, hypersensitive response) or abiotic stresses (like salt stress, water deprivation) (Figure S6). Several BPs were related to metabolic processes, more specifically to nucleotide metabolism and lipid metabolism and two terms were related to development (pollen development and leaf senescence). Shared mechanisms and gene families are summarized on Figure 5 also highlighting examples of genes found with highest A/S in several species.

**Figure 5.**
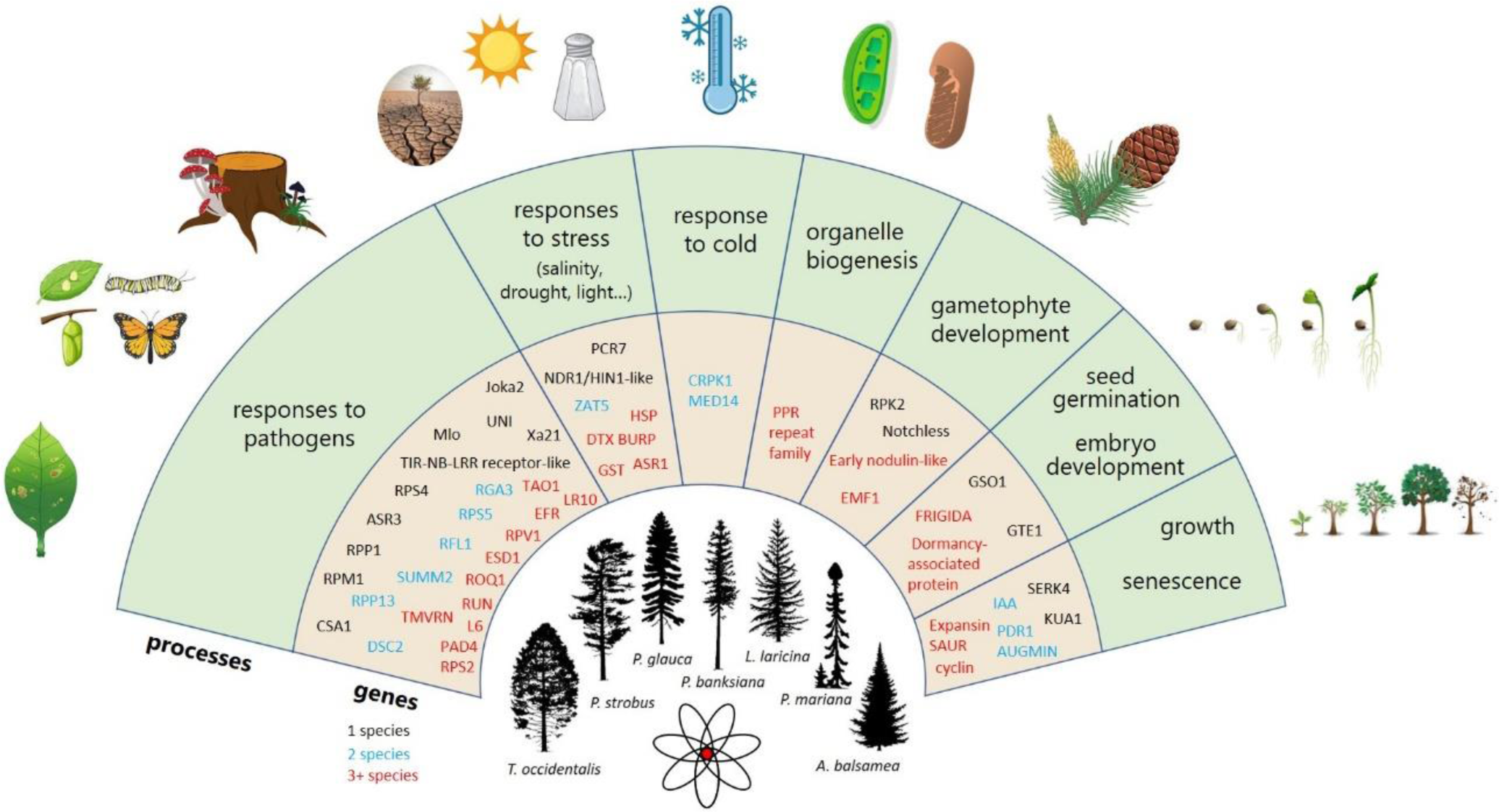
Summary of the conserved biological processes across the 300 genes with the highest A/S values for each of the seven conifer species analyzed. The outer circle shows the biological processes found with some degree of conservation across the seven species, the inner circle shows examples of genes in each functional category. Genes in black were found in a single species, those in blue were found in two species and those in red were found in three species or more. Source of images: https://tidcf.nrcan.gc.ca/;https://www.vecteezy.com/;https://www.freepik.com/;https://pixabay.com/

Genes with highest SNP A/S values were enriched in several GO classes, including 14 biological processes, 6 molecular functions and 2 cellular components (Figure 6). The BPs were summarized in five groups, the largest one representing responses to biotic stress and regulation processes. The enriched molecular functions were ADP binding, nucleotidase activity and protein binding (Figure 6). The two enriched cellular components were cytoplasm and chloroplast outer membrane (Figure 6). Several terms enriched in the annotation of these sequences were shared among the seven species (signal transduction, ADP binding, NAD+nucleotidase). Several of these terms correspond to the conserved mechanisms illustrated on Figure 5.

**Figure 6.**
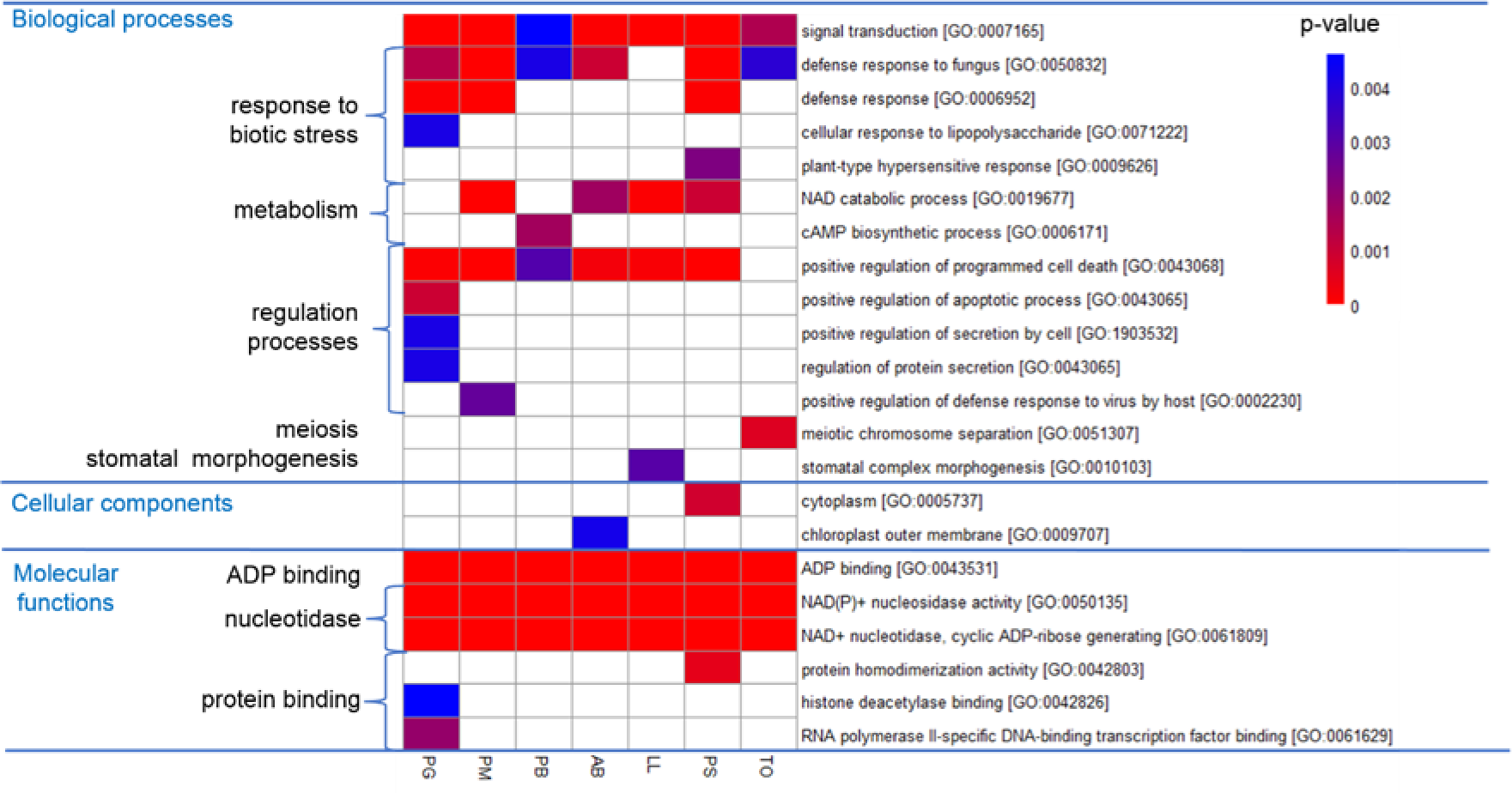
Gene Ontology terms enriched in the 300 genes with highest SNP A/S values for each of the seven conifer species analyzed, compared with the overall gene set for these species. (PG: *Picea glauca*, PM: *Picea mariana*, PB: *Pinus banksiana*, AB: *Abies balsamea*, LL: *Larix laricina*, PS: *Pinus strobus*, TO: *Thuya occidentalis*). The heatmap is based on the p-values of the enrichment tests (Fisher tests) and the color scale illustrates statistical significance. White cells represent non-significant tests at a threshold of 0.005. Summarized GO terms appear on the left.

Similar results were obtained when retaining the genes with top 600 SNP A/S values for each species (Figure S7), corresponding to lower minimum SNP A/S values of 1.87 to 2.34 depending of the species. Overall, the proportions of shared genes between species for these augmented groups of genes with large SNP A/S values were similar to that observed among the seven groups of 300 genes with top A/S values per species, whether grouping gene sequences by orthogroups, PFAMs or BPs (Figure S7).

### Gene sequences with an excess of nonsynonymous SNPs in conifers and homologous to positively selected genes in Brassica or poplar

Sequence similarity searches among conifer genes with highest SNP A/S values (genes with top 300 SNP A/S values per species) identified 340 homologs to positively selected genes (PSG) in poplar or *Brassica*, including 51 genes under positive selection in both poplar and *Brassica*. This set of 51 genes was partly redundant and matched a total of 20 distinct *Arabidopsis* genes including five transcription factors, four glutathione transferases, and a range of other families (Table 3).

**Table 3.** Accession number and description of 20 *Arabidopsis* gene sequences whose homologs are positively selected genes in at least one conifer species, and in poplar (Lin et al. 2018) and *Brassica* (Guo et al. 2017).

| Locus Identifier | Gene names | Protein family (known function) |
| --- | --- | --- |
| AT2G40270 |  | Protein kinase family protein |
| AT4G29270 |  | HAD superfamily, subfamily IIIB acid phosphatase |
| AT3G22142 |  | Protease inhibitor/seed storage/LTP family protein |
| AT5G03610 | GGL25 | GDSL-motif esterase/acyltransferase/lipase |
| AT4G15920 | AtSWEET17 | vacuolar fructose transporter (involved in control of leaf fructose content) |
| AT1G49200 | T ÑXICOS EN LEVADURA 75 (ATL75) | RING/U-box superfamily protein |
| AT3G14330 | CHLOROPLAST RNA EDITING FACTOR 3 (CREF3) | pentatricopeptide repeat protein (involved in chloroplast mRNA editing) |
| AT2G36800 | UDP-GLUCOSYL TRANSFERASE 73C5 (UGT73C5)<br>DON-GLUCOSYLTRANSFERASE 1 (DOGT1) | DON-Glucosyltransferase (presumably involved in the homeostasis of those steroid hormones) |
| AT1G78380 | GLUTATHIONE S-TRANSFERASE TAU 19 (GSTU19)<br>GLUTATHIONE TRANSFERASE 8 (GST8) | glutathione transferase from the Tau GST gene family (Expression is induced by drought stress, oxidative stress, and high doses of auxin and cytokinin) |
| AT5G15150 | HOMEBOX 3 (ATHB-3)<br>HOMEBOX FROM ARABIDOPSIS THALIANA 7 (HAT7) | homeobox-containing gene |
| AT5G40360 | MYB DOMAIN PROTEIN 115 (AtMYB115) | MYB family of transcription factors (involved in regulation of glucosinolate biosynthesis) |
| AT5G10280 | MYB DOMAIN PROTEIN 92 (ATMYB92) (ATMYB64) | MYB family of transcription factors |
| AT5G35550 | MYB DOMAIN PROTEIN 123 (AtMYB123)<br>TRANSPARENT TESTA 2 (TT2) | MYB family of transcription factors (acts as a key determinant in the proanthocyanidin accumulation of developing seed) |
| AT4G28530 | KIR1 (KIRA1)<br>NAC DOMAIN CONTAINING PROTEIN 74 (NAC074) | NAC family of transcription factors (regulates programmed cell death of stigmatic tissue) |
| AT5G05340 | PEROXIDASE 52 (PRX52) | peroxidases (involved in lignin biosynthesis) |
| AT1G17020 | SENESCENCE-RELATED GENE 1 (ATSRG1) | Fe(II)/ascorbate oxidase gene family (senescence-related gene) |
| AT3G25420 | SERINE CARBOXYPEPTIDASE-LIKE 21 (scpl21) | serine carboxypeptidase-like |
| AT4G34131 | UDP-GLUCOSYL TRANSFERASE 73B3 (UGT73B3) | UDP-glucosyl transferase |
| AT2G36750 | UDP-GLUCOSYL TRANSFERASE 73C1 (UGT73C1) | UDP-glucosyl transferase |
| AT3G46670 | UDP-GLUCOSYL TRANSFERASE 76E11 (UGT76E11) | UDP-glucosyl transferase |

The annotations of these 340 Angiosperm PSG were diverse but some of them could be clustered into groups of BPs (Figure S8). The analysis also identified a core set of 25 BPs shared between the Angiosperm and the seven conifer species, and identified BPs associated with genes with highest SNP A/S values in conifer species. The most represented processes in this group were responses to pathogens and oxidative stress, RNA modification, protein autophosphorylation and cell wall organization (Figure 7). Many sequences were homologous to disease resistance genes including several members of the RPS family conferring resistance against *Pseudomonas syringae*, homologs to SUMM2 (SUPPRESSOR OF MKK1 MKK2 2), LR10 (LEAF RUST 10 DISEASE-RESISTANCE), TMV resistance protein N, and MLO6. The disease resistance genes represented 19 families, four of which (RUN1, RPV1, ROQ1, TAO1) being also found in the sequences with highest SNP A/S values (Table S6). There were 44 genes homologous the RUN1 protein conferring resistance to mildew. Several gene families were involved in resistance against *Pseudomonas syringae* (RPS2, RPS5, RFL1, TAO1, RPP3, RPM1), many of them being also found among the homologs to positively selected genes identified in *Brassica* and poplar (Table S6).

**Figure 7.**
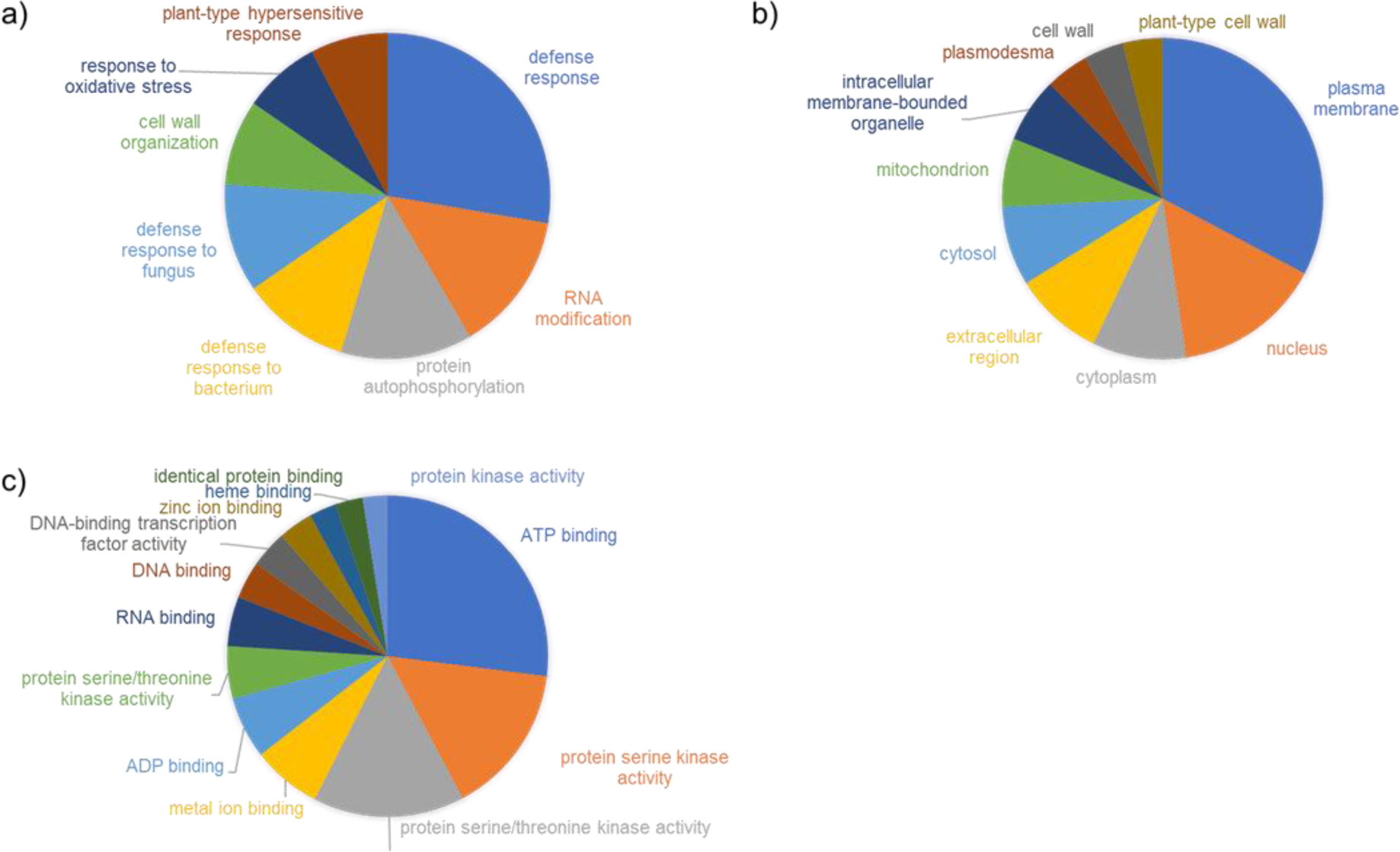
Most represented GO terms (found 10 times or more) in annotations of 340 conifer genes with highest SNP A/S values in the seven conifer species analyzed and found to be homologous (blastp E-value<1E-30) to posotovely selected genes (PSG) in poplar (Lin et al., 2018) or *Brassica* (Guo et al., 2017). a) biological processes, b) cellular components, c) molecular functions.

## DISCUSSION

### Contrasting levels of molecular genetic diversity among conifers

Several lines of evidence indicated that levels of total molecular genetic diversity differed substantially among the seven conifers (Table 1; Methods S7; Table S2). *Pinus strobus* and *Thuja occidentalis* had the lowest overall SNP diversity in their transcriptome, the two spruces were the most diverse, and the three remaining species were intermediate (Figure 1). Interpreting transcriptome-wide genetic diversity patterns is not straightforward because intraspecific variation results from the complex interplay between mutation rate, effective population size (long-term *N*e, which depends itself on historical and demographic factors), and linked selection (the molecular genetic diversity-reducing effect of selective sweeps on neutral loci in linkage with selected loci) (Ellegren & Galtier, 2016).

The seven species analyzed here are not expected to have significantly different mutation rates, as they are all long-lived woody perennials (Petit & Hampe, 2006; Sung et al., 2012), and considering that our genetic diversity estimates derive from transcriptome-wide SNP data rather than specific genes. However, in relation to the neutral theory of evolution which postulates that much of the standing genetic variation derives from neutral or nearly neutral mutations (e.g. Kimura, 1983; Ohta, 1992), part of the observed interspecific differences in molecular genetic diversity likely relates to historical effective population sizes (Bousquet et al., 1992). Indeed, the minimum historical population size (*N*e) of *Picea glauca* and *Picea mariana*, the most diverse species group in this study, was estimated at ∼100,000 or more individuals (Bouillé & Bousquet, 2005; Chen et al., 2010), while that of *Pinus strobus*, which belongs to the low diversity group, has been estimated to be one order of magnitude lower (5000 to 10,000 individuals; Zinck & Rajora, 2016). Likewise, *Thuya occidentalis*, the less genetically diverse species studied herein, also harbors a low *N*_e_ (Pandey & Rajora, 2012). There was also an apparent relation between the level of intraspecific SNP diversity and the size of species range (Figure 1), though the study of more species would be needed to confirm this trend. For instance, the two most diverse species have wide transcontinental distributions, while the two less diverse species are the most geographically restricted. As the number of glacial lineages of the Pleistocene era is usually positively related with range size in North American tree taxa (Jaramillo-Correa et al., 2009), widely-distributed species are likely to have retained larger historical population size and standing genetic variation than species with currently more restricted natural ranges.

The fact that molecular genetic diversity estimates reported herein originate from transcriptomic data makes it very likely that selection also played a role in shaping molecular genetic diversity, because the transcriptome mostly encodes functional information. Natural selection can constrain intraspecific molecular genetic diversity through hard sweeps (i.e when an arising beneficial mutation becomes rapidly fixed, leading to a reduction in neutral variation at linked sites; e.g. Smith & Haigh, 1974; Gillepsie, 2001) or soft sweeps (i.e. when multiple beneficial mutations at the same locus sweep simultaneously through the population (e.g. Smith & Haigh, 1974; Hermisson et al., 2005) to a lesser degree. Although selective sweeps are presumably uncommon in plants (Wright & Gaut, 2005; Grossman et al., 2010) including in conifers (Palme et al., 2008; Pavy et al., 2012b; Eckert et al., 2013), evidence for selective sweeps has been reported in some conifers (Eckert et al., 2009; Wang et al., 2020; Gagalova et al., 2022). Even if it was not possible in this study to test directly for the existence of such sweeps given the pool sequencing data obtained for each species studied herein, the number of synonymous SNPs and the SNP A/S ratio were negatively correlated; furthermore, genes with high SNP A/S values accumulated nonsynonymous SNPs and were virtually free of synonymous SNPs (Figure S1). This pattern is consistent with signatures of selective sweeps (Pavy et al., 2013; Garud et al., 2015), especially in forest trees where adaptation has been shown to be primarily polygenic (e.g. Le Corre & Kremer, 2012; Hornoy et al., 2015; Yeaman et al., 2016). This implies that various allelic combinations in different genes would yield the same phenotypic outcome via epistatic interactions (Le Corre & Kremer, 2012; Csilléry et al., 2018). Even with accumulating evidence of footprints of selective sweeps in conifers (e.g. Namroud et al., 2010; De La Torre et al., 2021; Gagalova et al., 2022), the occurrence of selective sweeps affecting much of the transcriptome remains hypothetical, as such signatures would remain hard to disentangle from effects of historical demographic fluctuations (Pavy et al., 2012b). In addition, it is unlikely that selective sweeps are a major determinant of transcriptome-wide genetic diversity in conifers because linkage disequilibrium decays rapidly within gene limits in conifer natural populations (e.g. Pavy et al., 2012b; De La Torre et al., 2017), therefore restricting the possible loss of neutral diversity surrounding selected loci. These observations suggest that, although both *N*_e_ and linked selection may have contributed to shaping molecular genetic diversity at the intraspecific level, the former is more likely to have been the main driver of differences in overall SNP diversity observed herein among the transcriptomes of the studied species.

### Relationship between overall SNP diversity and adaptive variation

The positive relationship observed between the overall SNP diversity of the transcriptome of each species and their proportions of genes with high SNP A/S values (Figure 2) suggests that standing genetic variation can constrain variation of more adaptive nature. It is well established that the efficiency of selection increases along with the effective population size, by reducing the strength of genetic drift and therefore limiting the loss of beneficial alleles as well as the fixation of deleterious ones (Charlesworth, 2009). Consistently, our results show that species of presumably larger historical population sizes carry the most adaptive variation. This is also in agreement with the idea that effective population size is the main determinant of local adaptation in plants (Leimu & Fischer, 2008), which has been reported for several species investigated herein, namely white spruce (Namroud et al., 2008; Hornoy et al., 2015; Depardieu et al., 2021), black spruce (Prunier et al., 2011;2012), jack pine (Cullingham et al., 2014), and eastern white pine (Nadeau et al., 2016). Because environmental adaptation is highly polygenic in conifers and involves heterogeneous gene responses (e.g. Hornoy et al., 2015; Yeaman et al., 2016), high standing genetic variation associated with large historical population size likely improves species adaptative potential by increasing the number of genetic trajectories to achieve adaptation. Hence, such flexibility may allow species to cope with a wider range of environmental conditions (i.e. gain the ability to colonize larger natural range and/or increase their ecological amplitude) and selective pressures (i.e. biotic and abiotic pressures encountered across their range).

### Extent of molecular and functional convergence among conifer adaptive genes

Despite the high overlap among gene sequences of the seven conifer species (Figure 3A), we only found limited molecular convergence among their sets of genes with highest SNP A/S values (Figure 3B; Table 2A). Convergence appeared equally limited at the gene family (PFAM) hierarchical level (Figure 3D), a result consistent with the pattern of species-specific expansion of large paralogous gene families in conifers (reviewed by De La Torre et al., 2020). The extent of molecular genetic convergence among taxa is expected to increase with their phylogenetic proximity, as a result of shared ancestry (Losos, 2011; Storz, 2016). However, no pattern related to phylogenetic relatedness among taxa was evident, with no sign of increased convergence among the two pairs of congeneric taxa that would have diverged the most recently (divergence between *Picea glauca* and *Picea mariana* ∼10 Mya; divergence between *Pinus strobus* and *Pinus banksiana* ∼85 Mya) (Table 2A). These observations suggest that each species followed a largely distinct adaptive path, and that adaptive convergence at the molecular genetic level appears to be limited in such reproductively isolated and phylogenetically distant conifers.

There is also strong evidence that the markerd divergence among species in their gene sets with high SNPS A/S values would be primarily driven by natural selection, rather than stochastic processes such as mutation or genetic drift (e.g. Losos, 2011; Mosca et al., 2012; Storz, 2016). The low levels of convergence observed among these gene sets either indicate that gene functional redundancy would allow species to cope with similar selective pressures using alternative genes, and/or that species experienced heterogeneous selective pressures throughout their historical and extent natural ranges. Our data support the first hypothesis, as functional convergence in biological processes among species gene sets with high SNP A/S values was quite higher than their limited overlap in gene content (Figure 3). This partly decoupled pattern indicates that the studied species would have sufficient metabolic and gene network flexibility to evolve alternative responses to the various selective pressures they faced under temperate and boreal climate regimes. It is also consistent with gene family expansions in conifers, implying some redundancy in gene functions (Guillet-Claude et al., 2004; Bedon et al. 2010; Pavy et al., 2012a; Stival Sena et al. 2018; Van Ghelder et al. 2019). This redundancy at the functional level may have had significant evolutionary implications for the persistence of these northern conifer species during millions of years, in the face of geological climate instability and in spite of demographic fluctuations. For instance, with the multiple glaciation cycles of the Pleistocene era in eastern North America, signatures of demographic fluctuations such as bottleneck effects or founder events have been detected in various conifer species from northeastern North America, which would have implied more or less important losses of overall genetic diversity (Perron et al., 2000; Gamache et al. 2003; Godbout et al., 2010; Namroud et al., 2010).

Noticeably, while the Cupressaceae *Thuja occidentalis* had the lowest overall SNP diversity in its transcriptome (Table 1), it carried the highest number of species-specific genes with highest SNP A/S values among the seven studied conifers (Figure 3F), which is likely the consequence of its more phylogenetically remote position from all other Pinaceae taxa. These *Thuja occidentalis* genes were also representative of the highest number of BPs among the conifers studied, resulting in the lowest number of genes per BP (Table 2B). One can hypothesize that gene redundancy is likely lower in *Thuja occidentalis*, because it harbored both the smallest number of genes per BP and, at the same time, the smallest proportion of genes with high SNP A/S values (Figure 2).

In addition to the gene redundancy hypothesis, it is possible that the low convergence observed among conifer taxa in genes with high SNP A/S values also reflects the fact that they cope with species-specific selective pressures throughout their large natural ranges. Some species such as the two *Picea* spp. have large ecological amplitude and a transcontinental range across which they may encounter a variety of selective pressures related to biotic and abiotic stresses (Nienstaedt & Zasada, 1990; Viereck & Johnston, 1990). Also, *Picea glauca* would prefer mesic sites (Hornoy et al., 2015) while *Picea mariana* could adapt to a larger variety of site conditions (Lo et al., 2023). Others have more distinct preferred habitats, such as wet sites for *Larix laricina* (Cheliak et al., 1988), dry sites for *Pinus banksiana* (Rudolph & Laidly, 1990), or both for the more extremophile *Thuja occidentalis* (Matthes-Sears & Larson, 1991), which can trigger specific adaptive responses and explain the finite extent of convergence among the species studied.

Hence taken together, our results suggest that the adaptive trajectories of these conifer species were likely shaped by the interplay of gene redundancy and heterogeneous selection landscape, and that these two drivers likely contributed to the low convergence observed in terms of gene sets displaying high SNP A/S values, and higher convergence at the molecular functional level.

This pattern aligns well with other reports of low estimates of molecular genetic convergence among conifers. A study of adaptive traits in four alpine conifers from the *Pinus*, *Abies* and *Larix* genera identified only seven climate-associated genes shared by two or more species out of several hundreds of sequences analyzed (Mosca et al., 2012). Another study uncovered only 47 convergent genes (representing between 10% and 18% of all genes putatively under selection) involved in local adaptation in a spruce and a pine taxon that diverged ∼150 Mya (Yeaman et al., 2016). Also, a transcriptome-wide survey of genetic variation in the two quite closely-related but ecologically contrasted *Picea glauca* and the coastal *Picea sitchensis* from the Pacific Northwest, revealed only 15 shared genes out of hundreds of genes showing selection footprints (Gagalova et al., 2022). Similar modest molecular genetic convergence was also reported in Angiosperms such as in *Arabidopsis* (Guggisberg et al., 2018; Preite et al., 2019), or among taxa from the Brassicaceae family (Rellstab et al., 2020).

### Nature of molecular functional convergence among conifers and between conifers and Angiosperms

Functional annotations of genes with high SNP A/S values were associated with a great variety of molecular functions and biological processes (Figure S2), in agreement with the polygenic nature of adaptive traits and the many empirical studies reporting a wide range of genes and functions underlying them in conifers (e.g. Prunier et al., 2011; Mosca et al., 2012; Hornoy et al. 2015; Depardieu et al., 2021).

Despite this large functional diversity, the core set of functions and processes shared by all conifer species analyzed revealed a clear pattern of shared adaptive evolution at the functional level. Indeed, half of the 31 shared BPs were related to environmental stress responses, and mechanisms related to defense against pathogens (responses to biotic stress, programmed cell death) were widely represented and enriched in rapidly evolving genes (Figure 5; Figure 6). We found many homologs of disease resistance genes with the NB-ARC domain (Figure S4) and several gene families involved in resistance to *Xanthomonas*, *Pseudomonas* or rusts (Table S6). This indicates that selection pressures exerted by pathogens are likely ubiquitous in conifers and play a prominent role in their adaptation to environment. Similarly, several shared BPs were linked to abiotic stress response, and more specifically to water and oxygen stimuli (i.e. cellular response to hypoxia, response to oxidative stress, or response to water deprivation), suggesting that drought and flooding could also be drivers of adaptive evolution at the functional level in conifers.

Mechanisms with indirect but crucial roles in stress response such as RNA modification and regulatory mechanisms were also found in sequences with highest SNP A/S values (Figure 5). Regarding RNA modification, we identified 27 sequences encoding pentatricopeptide repeat (PPR) proteins, among which two were also reported as convergent adaptive genes in pine and spruce taxa (Figure S4; Yeaman et al., 2016). The PPR proteins have fundamental roles in organelle biogenesis and function, being involved in photosynthesis, respiration, development and environmental responses (Barkan & Small, 2014). Positive regulation of programmed cell death, a process known to be involved in response to biotic and abiotic stresses in plants (60), was also enriched in all Pinaceae species (Figure 6). Sequences encoding transcriptional regulators included several MYB and WRKY sequences, and a homolog of the Scarecrow-like p9 transcription factor involved in plant development in *Arabidopsis* (accession SCL9_ARATH). We also found a homolog of KUA1 (accession KUA1_ARATH; Figure 5), a transcriptional repressor of several genes involved in chloroplast function and responses to light and auxin, which regulates leaf growth, development and senescence (Lu et al., 2014; Huang et al., 2015). Homologs of genes involved in seasonal transitions in *Arabidopsis* are likely prime targets of natural selection given that they contribute to the adaptation of plants to their environment, assuming that their functions are conserved across seed plants. Key examples include EMF1 (EMBRYONIC FLOWER1; accession EMF1_ARATH) and a FRIGIDA-like protein (accession FRL3_ARATH), which both regulate phase transition during shoot, flower and seed development (Figure 5). In *Arabidopsis,* the transcriptional repressor EMF1 controls vegetative development by delaying the transition to reproductive phase and the initiation of flowering and is involved in response to salt stress (Pu et al., 2013), whereas FRIGIDA sequences are required for the winter-annual habit (Michaels et al., 2004) (Figure 5). The identification of several homologous regulators involved in survival in Angiosperms suggests a possible key role in conifers and makes them a relevant class of genes to target in future molecular biology studies.

We uncovered a shared set of 20 homologous genes putatively under positive selection in conifers and two Angiosperm model taxa (Brassicaceae and poplar). This level of molecular convergence was higher than expected, given that Angiosperms and Gymnosperms (to which belong conifers) diverged ∼350 Mya (Li et al., 2019). Disease resistance genes against pathogens are known to evolve rapidly in flowering plants (Meyers et al., 2005). Although well represented among genes with highest SNP A/S values in conifers, they were not predominant among the core set of convergent genes between conifers and Angiosperms. In contrast, several PPR genes involved in RNA editing, and several transcription factors such as MYBs showed up in this set of genes (Table 3). To our knowledge, such a molecular signature has not been reported to date and may be interpreted as a sign that adaptive convergence at the molecular level can take place at a broad taxonomic level. Consequently, these genes represent valuable candidates for future evolutionary studies aiming to characterize molecular and functional convergence among seed plants.

## CONCLUSIONS

Transcriptome-wide SNP diversity was assessed for seven partially sympatric and reproductively isolated conifers. We found marked variation in overall SNP diversity among species, that would reflect differences in demography and historical population size. Little overlap in sets of adaptive genes under positive selection was noted between species, suggesting distinct evolutionary trajectories. In contrast, their biological functions were much convergent and largely related to stress response and regulatory mechanisms. This trend indicates high molecular plasticity in response to similar climate and natural selective pressures. A number of adaptive genes was found shared between conifers and Angiosperms, despite their ancient divergence ∼350 Mya.

## EXPERIMENTAL PROCEDURES

### Biological materials

Seven conifer species were sampled, namely *Picea glauca*, *Picea mariana*, *Pinus strobus*, *Pinus banksiana*, *Abies balsamea*, *Larix laricina*, and *Thuja occidentalis*. Seeds from ten provenances were obtained from the National Tree Seed Center (Fredericton, New-Brunswick, Canada), paying special attention to avoid provenances located within sympatric or paratric zones in species known to spontaneously hybridize with related taxa (Figure 1; Methods S1). For each species, twenty seeds from ten distinct provenances per species (two seeds per provenance) were dissected to extract the embryos, which were flash frozen in separate tubes. To identify and exclude paralogous non-mendelian SNPs, four provenances were randomly selected per species and one seed per provenance was dissected to extract the haploid megagametophyte, which was flash frozen in liquid nitrogen in separate tubes. In total, four haploid megagametophytes and between 15 and 18 diploid embryos were extracted and flash frozen for each species, prior to library synthesis (Methods S1).

### Sequencing

Total RNA was extracted using the MasterPure™ Plant RNA Purification kit (Epicenter, Madison, WI, USA). RNAs were sequenced in paired-end mode (2×125 bp) with an Illumina HiSeq 2500 (Methods S2). Raw sequencing data (reads) were deposited in the public database ENA (European Nucleotide Archive, https://www.ebi.ac.uk/ena/browser/home, accessions ERS16017105-ERS16017139 and ERS16049778-ERS16049791) and vcf files containing variants identified in each species were deposited in DRYAD (https://datadryad.org/stash, DOI pending). We assessed the good representativity of the analyzed transcriptomes based on sequence similarity searches (Methods S3).

### SNP calling

After sequence quality controls and filtering (Methods S4), reads were aligned to the reference transcriptomes of each species previously published (Van Ghelder et al., 2019). SNPs were called using HaplotypeCaller, from the GATK tool kit (McKenna et al., 2008; DePristo et al., 2010) and subsequently quality-filtered (Methods S5). Since the megagametophyte is a haploid tissue in all seven conifer species investigated herein, SNPs identified within megagametophyte libraries were likely indicative of variations between paralogous gene sequences also occurring in embryos. Thus, these SNPs were considered as false-positives and were subtracted from SNPs identified in pools of embryos before subsequent analyses were carried out. In total, above 1.4 million raw SNPs were identified (Methods S5, Table S5.1). After removal of paralogous SNPs identified in haploid megagametophytes, more than 866K SNPs remained (Table S1). Among them, ∼398K high-quality SNPs were located in coding sequences representing almost ∼97K Open Reading Frames in total (∼16K transcripts per species) (Table S1). All species considered, ∼82% of the transcripts and ∼70% of the ORFs carried SNPs (Table S1).

Transcripts that were included in the comparison of molecular genetic diversity across species and the analysis of the functional annotations of polymorphic sequences had to present an average coverage of 10 reads or more, at least 300 nucleotides with a depth of 10 or more reads, and contain coding sequences.

### Estimation of SNP abundance in transcripts

The length and read depth of transcripts were heterogenous across the seven species investigated (Methods S6). It is essential to control for such effects before analyzing SNP abundance disparities across species. In this purpose, we applied a regression model assuming that the number of SNPs within transcripts follows a negative binomial distribution (Eo & DeWoody, 2012). The model corrected efficiently for variations among transcripts depth and length, thus enabling a rigorous comparison of the SNP diversity across species (Methods S7). SNP rate heterogeneity among species was tested using a Kruskall-Wallis test. To group species based on their level of total SNP diversity, Kolmogorov-Smirnov and Cramer-von Mises tests were conducted.

### Estimation of gene SNP A/S ratios

Based on the premise that nonsynonymous mutations are predicted to contribute more to phenotypic evolution than synonymous mutations (Stern & Orgogozo, 2008), one way to study molecular convergence is to compare the ratio of substitution rates at nonsynonymous (*K*a) versus synonymous (*K*s) sites in orthologous protein-coding sequences among species. Similar inferences can be drawn within taxa from gene SNP A/S ratios, since both ratios have been shown to be strongly positively correlated (Liu et al., 2008).

The SNP A/S ratio was calculated for each gene as the number of SNPs per nonsynonymous site (A) divided by the number of SNPs per synonymous site (S). An adjusted SNP A/S ratio was used to include genes with no synonymous SNPs following the empirical logit principle (Agresti, 2013):

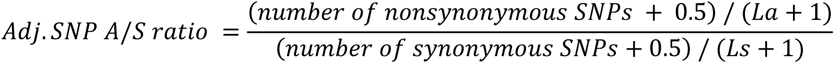

The SNP A/S ratio was calculated over the longest open reading frame predicted for each transcript. The SNP A/S ratio was weakly but significantly correlated to transcript coverage. Therefore, we had to apply an adjustment to neutralize this effect. The procedure correcting for both transcript length and depth is detailed in the Supporting information (Methods S8). This step was essential to enable a comparison of the sequences with the highest A/S values among species.

Comparison of genes exhibiting the highest SNP A/S values among species was carried out for a predetermined and equivalent number of genes in all species, which ensured representativity and comparability. Considering the large numbers of genes with SNP A/S values above 1, and even above 2 (Table S3), we opted to analyze the 300 genes with highest SNP A/S values in each species, which represented between 2% and 3% of the ORFs carrying SNPs and corresponded to A/S ratio threshold ranging between 2.51 and 2.88 (Table S3). This dataset was used to analyze and compare the sequences and functional annotations of these deemed fastest-evolving genes across the seven species included in the study. The analysis was also replicated with the 600 sequences with highest SNP A/S values (corresponding to A/S ratio threshold ranging between 1.87 and 2.34, which led to the same conclusions regarding molecular genetic divergence, and molecular functional convergence (Figure S7).

### Sequence annotation and analyses

Predicted protein sequences were clustered into orthogroups with OrthoFinder v2.3.8 (Emms & Kelly, 2019) run with default settings. Functional annotations of ORFs were derived from sequence similarity searches conducted with blastp against Uniprot (E-value <e^−15^) and PFAM (El-Gebali et al., 2019). Sequences were also assigned to Gene Ontology (GO) classes by using the mapping between the UniprotKB sequences and the GO terms. The heatmaps were generated using the pheatmap R package (Kolde, 2019).

Enrichment tests were conducted with the R package topGO (Alexa et al., 2006; https://bioconductor.org/packages/release/bioc/html/topGO.html), in order to identify GO terms enriched among annotations of the genes with the highest SNP A/S values (Methods S9).

### Genes with highest SNP A/S values in conifers and with positively selected homologs in Brassica or poplar

Positively selected genes (PSG) were identified in *Brassica* (Guo et al., 2017). Their corresponding *Arabidopsis* sequences were retrieved (https://www.arabidopsis.org/) for a total of 621 sequences. PSGs were identified in poplar (Lin et al., 2018). The *Populus trichocarpa* sequences were retrieved from PopGenIE.org. The 2,100 conifer sequences with highest SNP A/S values were compared at the protein level to sequences from poplar and from *Arabidopsis*, a species closely related to *Brassicas*. Overall, pairs of homologous sequences between these dicots and conifers were identified following a blastp search (E-value<1E-30). When one dicot gene sequence was found homologous to several conifer gene sequences, or when one conifer gene sequence was homologous to several dicot gene sequences, only the best match was selected.

## AUTHOR CONTRIBUTIONS

JB and JM designed the study and obtained the funding. NP and SG performed data analysis. JL, BB, PR, JP produced the sequence data and contributed to study design. GD conducted statistical analyses. JB, NP, SG wrote the paper.

## ACKNOWLEDGMENTS

This research was part of the GenAC project supported by a research grant from the Québec Ministry for the Economy, Science and Innovation to JM, JB and PR. We are grateful to Aida Azaiez, France Gagnon and Isabelle Giguère (Canada Research Chair in Forest Genomics, Univ. Laval) for providing technical assistance, as well as Jean Beaulieu (CRC Forest Genomics, Univ. Laval) for his valuable insights into early statistical analyses.

## DATA AVAILABILITY STATEMENT

All data generated or analyzed in this study will be made available in public repositories.

